# Fluorescence lifetime imaging as an *in situ* and label-free readout for the chemical composition of lignin

**DOI:** 10.1101/2021.08.26.457748

**Authors:** Sacha Escamez, Christine Terryn, Madhavi Latha Gandla, Zakiya Yassin, Gerhard Scheepers, Torgny Näsholm, Ola Sundman, Leif J. Jönsson, Judith Lundberg-Felten, Hannele Tuominen, Totte Niittylä, Gabriel Paës

## Abstract

Important structures and functions within living organisms rely on naturally fluorescent polymeric molecules such as collagen, keratin, elastin, resilin, or lignin. Theoretical physics predict that fluorescence lifetime of these polymers is related to their chemical composition. We verified this prediction for lignin, a major structural element in plant cell walls and one of the most abundant components of wood. Lignin is composed of different types of phenylpropanoid units, and its composition affects its properties, biological functions, and the utilization of wood biomass. We carried out fluorescence lifetime imaging microscopy (FLIM) measurements of wood cell wall lignin in a population of 90 hybrid aspen trees genetically engineered to display differences in cell wall chemistry and structure. We also measured wood cell wall composition by classical analytical methods in the wood cell walls of these trees. Using statistical modelling and machine learning algorithms, we identified parameters of fluorescence lifetime that predict the content of S-type and G-type lignin units, the two main types of units in the lignin of angiosperm plants. Finally, we show how quantitative measurements of lignin chemical composition by FLIM can reveal the dynamics of lignin biosynthesis in two different biological contexts, including *in vivo* while lignin is being synthesized in the walls of living cells.

## Introduction

Woody biomass represents the majority of the biomass on land^1^, and this abundance gives it an ecological importance such that it shapes evolution, as well as makes it a reservoir of renewable resources for a sustainable economy. This biomass mostly consists of specialized plant cell walls often referred to as lignocellulosic cell walls, on account that they consist mainly of three types of polymers: cellulose, hemicelluloses and lignin^2-5^.

Lignin is a particularly interesting component of lignocellulosic cell walls from both a biological and an economic point of view. Biologically, lignin plays a role in processes that rely on specific cell wall properties, such as preventing diffusion of solutes and pathogens through the walls^6-11^, providing greater rigidity and mechanical strength^12,13^, or allowing for clear-cut cell wall breaks when required, for example in organ abscission^8,14^. For biorefinery of lignocellulosic biomass, lignin is seen as both a potential source of high-value compounds^15,16^, as well as one of the major recalcitrance markers to the necessary fractionation of cell wall polymers for different applications^17-19^. These different, sometimes seemingly opposite properties of lignin arise from the fact that the lignin polymer is particularly complex.

Lignin is produced by polymerization in the cell walls of different types of monomers, mainly hydroxycinnamyl alcohols, which, once integrated as phenylpropanoids into the polymer, are known as G-type, S-type, and, to a lesser extent, H-type lignin units^2,13,20^. In the wood of leaved trees (Angiosperms), the majority of the lignin is composed of S-type and G-type units, while lignin can also include atypical units^20^. In addition to consisting of diverse units, lignin is also made structurally complex by the fact that a same pair of units can form different combinations of covalent bonds between one-another^2,13,20^. The unit composition, as well as the diversity of bonds between units, thus confer different properties to lignin^2,13,20^, which in turn influences the biological role a particular type of lignin can play, for example by providing a range of possible mechanical properties to the cell walls^21^.

Interestingly, owing to the molecular structures formed during polymerization, lignin is naturally fluorescent^22-24^. In theory fluorescence can be leveraged to measure the chemical composition of lignin *in situ*, with single cell wall resolution^25^. Several attempts at linking lignin chemistry to fluorescence have relied on monitoring the emission spectrum of lignin fluorescence^22,23,25^. These spectral differences in lignin fluorescence have, however, so far not been shown to correspond to specific lignin chemical compositions. One major hinder of such spectral analyses may reside in the fact that the detection of emission spectra of fluorophores can be strongly influenced by fluorophore type, interactions and concentration as well as external factors^26^. Another dimension of fluorescence, one that could be correlated more reliably with fluorophore chemical composition, should be leveraged in order to best exploit the potential of lignin fluorescence as *in situ* and label-free chemotyping method.

In line with the theoretical physics that explains atom fluorescence^27^, Strickler and Berg^28^ derived the equation that provides an account of the fluorescence lifetime of molecules, i.e. the time needed for a fluorophore to emit a photon after it has been excited. This so called Strickler-Berg equation predicts that fluorescence lifetime mostly depends on the chemical composition of the fluorophore, while remaining invariant to many of the factors that affect the emission spectrum, in particular concentration^26^. The potential of monitoring fluorescence lifetime has more recently been unlocked thanks to the development of advanced pulsed lasers^29^ allowing the measurement of fluorescence lifetime between two laser pulses, and by recent advances in hardware and methods for photon detection at the nanosecond time-scale^29,30^.

Recent studies have started to investigate the potential of using fluorescence lifetime to analyse lignocellulosic cell walls^22^. A model plant *Arabidopsis thaliana* genotype known to display altered lignin composition also showed differences in fluorescence lifetime compared to wild-type plants^31^. Furthermore, chemical treatment known to affect lignin chemical environment resulted in lignin fluorescent lifetime shifts that correlated with treatment severity on the lignocellulosic biomass of several plant species^32,33^. However, these observations of correlations between lignin fluorescence lifetime and lignin composition did not provide the parameters that would link specific changes in lignin composition with variations in fluorescence lifetime.

In this study, we therefore set out to develop fluorescence lifetime as a readout for lignin chemical composition. To this end, we assembled a population of genetically engineered trees with different wood cell wall properties, such as altered lignin unit composition. By monitoring wood cell wall chemical composition traits as well as recording parameters of fluorescence lifetime on these trees, we identified fluorescence lifetime parameters that can predict chemical composition traits. We also provided examples of how this new chemotyping method based on fluorescence lifetime imaging microscopy (FLIM) can be used to answer questions of biological and applied relevance. Finally, we discussed how FLIM could be used for in situ chemical composition analyses of other naturally fluorescent polymers.

## Results

### Chemically different lignocellulosic cell walls display different fluorescence lifetimes

As a proof of concept for the correlation between lignin fluorescence lifetime and the chemical composition of lignocellulosic cell walls, we performed fluorescence lifetime imaging microscopy (FLIM) on xylem tissues in stem cross-sections of *Arabidopsis thaliana* wild-types and mutants previously shown to display modified cell wall chemistry (Fig. 1).

**Figure 1.**
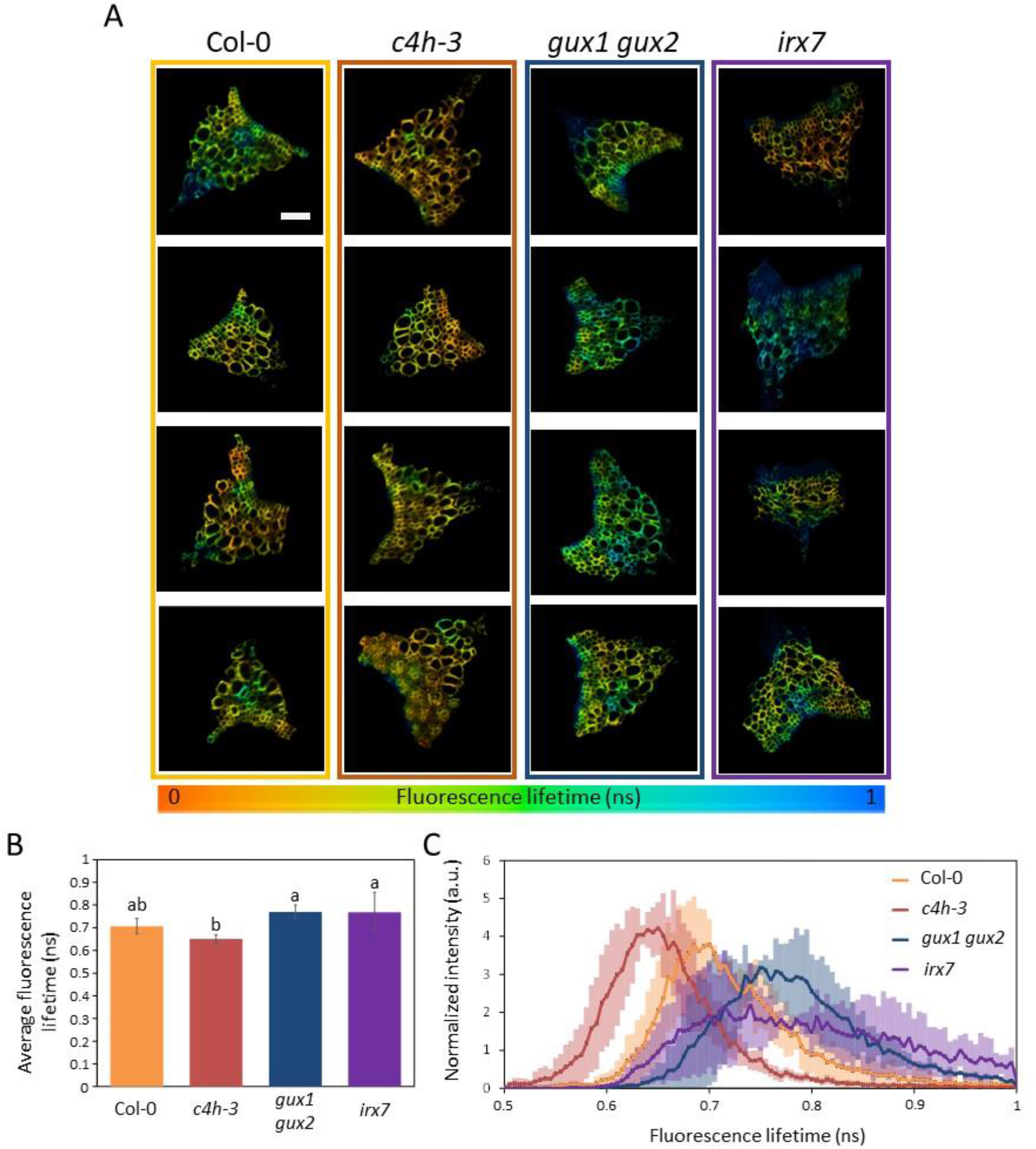
Fluorescence Lifetime Imaging Microscopy (FLIM) of *Arabidospsis* genotypes with different lignocellulosic cell wall properties reveals the potential of FLIM to image cell wall chemistry *in situ*. (A) Fluorescence lifetime imaging micrographs of a xylem bundle in *Arabidopsis thaliana* stem cross sections for the indicated genotypes. Xylem cell wall autofluorescence is color-coded based on average fluorescence lifetime for each pixel. Scale bar = 20 µm. (B) Average xylem cell wall fluorescence lifetime for each *Arabidopsis* genotype. Error bars represent standard deviation (n = 4 biological replicates), and genotypes that do not share any letter are significantly different according to a post-ANOVA Fisher’s LSD test. (C) Average distribution of xylem cell wall fluorescence lifetime for each genotype. Semi-transparent bars represent standard deviation (n = 4 biological replicates).

The *c4h-3* mutation targets an enzyme in the lignin biosynthetic pathway^34^, and had previously been described as triggering an increased ratio of S-type to G-type lignin^35^, the two main types of lignin in angiosperms, originating from two distinct monomers (Boerjan et al., 2003). In line with their altered lignin chemical composition, the *c4h-3* mutants showed a down-shift in lignin fluorescence lifetime profile compared to that of *Col-0* wild-type plants (Fig.1A-C). The *gux1gux2* double mutant is unable to add glucuronic acid and 4-O-methylglucuronic acid (collectively referred to as MeGlcAc) side groups on xylan backbones of hemicelluloses^36,37^. These MeGlcAc groups of xylan hemicelluloses are thought to covalently associate with lignin^38^. Their absence in the *gux1gux2* mutant should theoretically result in altered lignin fluorescence lifetime: this is confirmed by a shift towards higher lifetime values in *gux1gux2* mutants compared with wild-type (Fig. 1A-C). Finally, the *irx7* mutant is defective in biosynthesis of xylan hemicelluloses^39^, which affects the overall cell wall matrix, and, likely indirectly, lignin and its fluorescence lifetime (Fig. 1A-C).

Notably, the genotypes differed not only in terms of average fluorescent lifetime with statistical significance (Fig. 1B), but more evidently in the distribution of the observed lifetimes (Fig. 1C), with large differences in the maximum wavelength values. In other words, parameters of the fluorescence lifetime appeared as variables that could predict aspects of the lignocellulosic cell walls chemical composition. We therefore set to explore which fluorescence lifetime parameters could be used to predict lignocellulosic cell walls’ chemistry.

### Genetically engineered trees for cell wall chemistry as standards for FLIM as a chemical readout

To look for predictive correlations between lignocellulosic cell wall fluorescence lifetime and chemical composition, we assembled a collection of hybrid aspen trees (*Populus tremula x tremuloides*) genetically engineered for modified cell wall properties (Dataset S1). Using *Populus* trees as a model is justified by the fact that *Populus* species (poplars, aspens, and hybrids) have been established as model tree species^40^, with several members having sequenced genomes^41-43^, and *Populus* trees hold great potential as feedstocks for wood biomass^44,45^.

To further increase the variation in cell wall properties, we also submitted these *Populus* genotypes to two different treatments, consisting in having the majority of their nitrogen nutrition from a mineral or an organic source (Dataset S2). Previously, different amounts of nitrogen fertilization had been shown to alter lignin accumulation in the wood of hybrid aspens^46^. In another *Populus* hybrid species, differential nitrogen fertilization resulted in alterations of wood lignin composition and structure^47^. We therefore reasoned that different sources of nitrogen nutrition (with organic nitrogen bearing similarities to low nitrogen fertilization in terms of tree physiology^48^) may add variability to the lignin properties such as abundance of different lignin units, or to overall lignin accumulation in the wood of our hybrid aspen collection.

In the resulting collection of 90 trees (11 genotypes with two nutrition treatments, including each 4-5 replicates per genotype and per treatment), we measured 10 traits of the chemical composition of the lignocellulosic cell walls, such as the carbohydrate content (combining cellulose and hemicelluloses), total lignin content, as well as the proportions of cell wall lignin made of S-type, G-type and H-type units (Fig. 2A; Dataset S3). Furthermore, we measured the release of five different monosaccharides after enzymatic hydrolysis of the cell walls, so called saccharification, either without or after a potent acidic pretreatment (Fig. 2B; Dataset S3). Finally, we performed FLIM imaging on stem cross-sections of the 90 trees (Figure S1) to register 18 parameters of fluorescence lifetime (Fig. 2C; Dataset S3).

**Figure 2.**
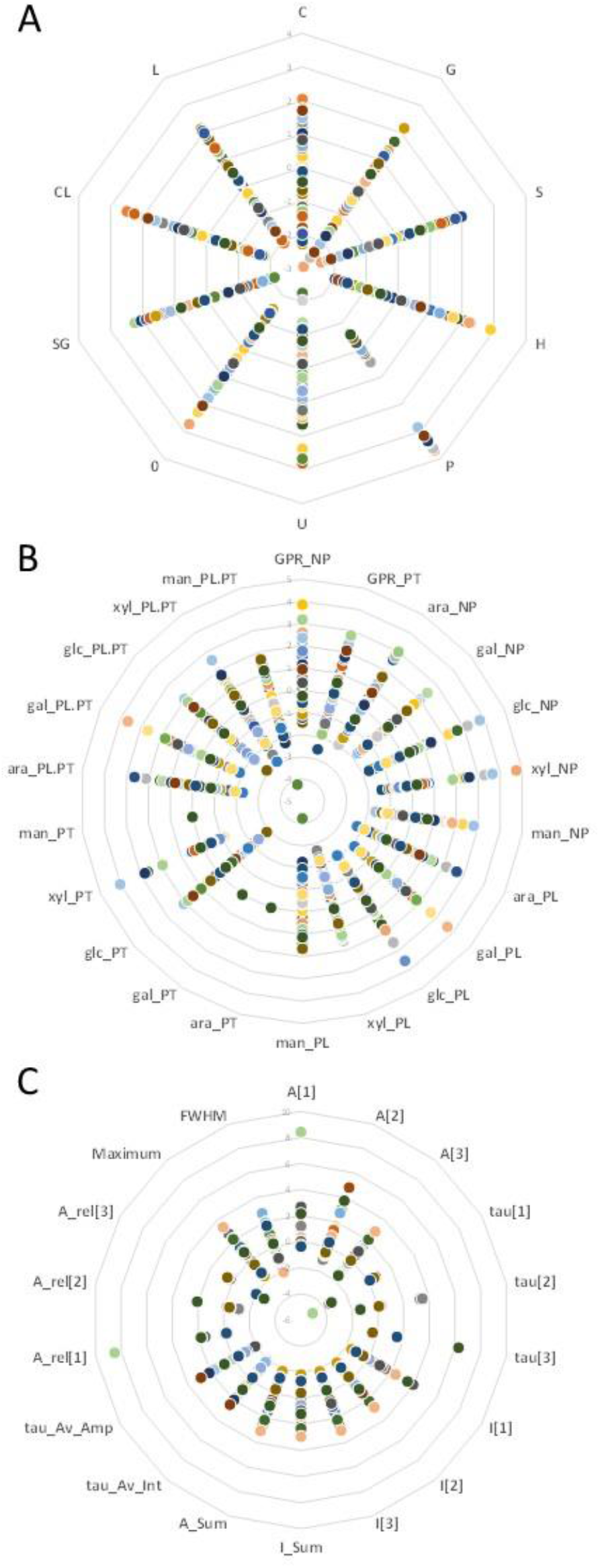
The collection of 90 hybrid aspens displays variation in FLIM parameters, chemical composition and saccharification, allowing for statistical modelling. (A) Variation in cell wall chemical composition traits (scaled and centered) measured by pyrolysis-GC/MS. Labels indicate the wood content in C: carbohydrates; L: total lignin; G: G-type lignin; S: S-type lignin; H: H-type lignin; P: other phenolic lignin units; U: unidentified compounds; 0: unknown compounds; SG: S-type/G-type lignin ratio; CL: carbohydrate/lignin ratio. (B) Variation in sugar released from saccharification (scaled and centered) without (NP) or after pretreatment (in pretreatment liquid PL and in hydrolysate PT). Labels indicate GPR: glucose production rate; ara: arabinose; gal: galactose; glc: glucose; xyl: xylose; man: mannose. (C) Variation in FLIM parameters (scaled and centered). The average lifetime of each image was best described as resulting from three distinct lifetimes (3-exponential fit). Labels indicate A[n]: amplitude for lifetime [n]; tau[n]: average lifetime value for lifetime [n]; I[n]: intensity for lifetime [n]; I_Sum: sum of intensities considering all three lifetimes; A_Sum: sum of amplitudes considering all three lifetimes; tau_Av_Int: average lifetime value; tau_Av_Amp: amplitude of the average lifetime. A_rel[n]: relative contribution of lifetime [n] to the average lifetime.

Nearly all of the traits showed a great variation, with values covering rather evenly the ranges between the mean (set to 0 in scaled and centred values) and the extremes (Fig. 2A,C). This variation, and the fact that it is distributed across the full range of values, enables using the measured traits as variables in multivariate analyses and statistical modelling.

### Fluorescence lifetime parameters accurately predict G-type and S-type lignin content

As expected from the wide range of variations in cell wall chemical composition in the *Populus* collection, differences in lignin fluorescence lifetime could be detected between genotypes (Fig. 3A,B; Fig. S1). Fluorescence lifetime differences could even be seen between cell walls of different wood cell types (Fig. 3A,B), suggesting that FLIM could be used to detect differences in chemical composition *in situ* if lifetime parameters could accurately predict chemical composition traits.

**Figure 3.**
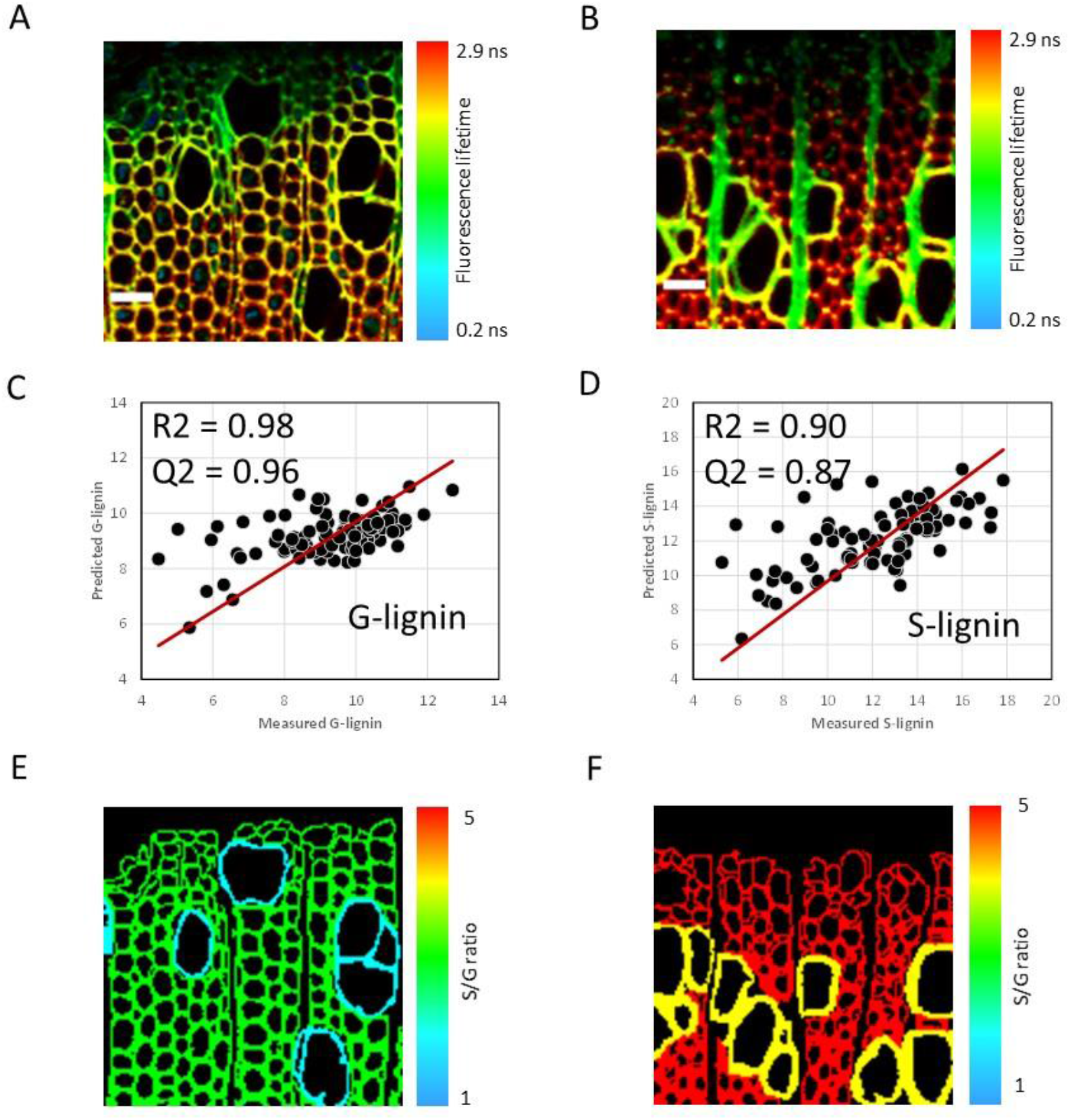
A set of fluorescence lifetime parameters predicts lignocellulosic cell wall S-lignin and G-lignin content *in situ*. (A) Fluorescence lifetime imaging micrographs of wild-type developing wood (directly next to the cambium, the non-fluorescent area at the top of the image). Scale bar = 30 µm. (B) Fluorescence lifetime imaging micrographs of *c4h*-RNAi (2081-B) developing wood section (directly next to the cambium, the non-fluorescent area at the top of the image). Scale bar = 30 µm. (C) Scatter plot of the G-lignin content, measured versus predicted with the best statistical model (Dataset S4). R2 indicates the fraction of the variance that is explained by the model, while Q2 evaluates the predictivity of the model (1 ≤ Q2 < -∞, with 1 representing a perfect prediction). (D) Scatter plot of the G-lignin content, measured versus predicted with the best statistical model (Dataset S4). R2 indicates the fraction of the variance that is explained by the model, while Q2 evaluates the predictivity of the model (1 ≤ Q2 < -∞, with 1 representing a perfect prediction). (E) Micrograph of wild-type developing wood (from A), where the vessels on the one hand, and the fibres on the other hands, were pseudo-colored based on the average S/G ratio for this cell type, as predicted in each sample by the statistical models for S-type and G-type lignin content (C, D, and Dataset S4). (F) Micrograph of *c4h*-RNAi (2081-B) developing wood (from B), where the vessels on the one hand, and the fibres on the other hands, were pseudo-colored based on the average S/G ratio for this cell type, as predicted in each sample by the statistical models for S-type and G-type lignin content (C, D, and Dataset S4).

To predict cell wall chemical composition, or even saccharification traits, based on fluorescence lifetime parameters, we employed statistical modelling following an established methodology^18,43^. Briefly, we employed three types of models: linear models, or the more complex machine learning algorithms known as generalized additive models^49^ and Random Forests^50^. For each cell wall chemical composition trait as well as a selection of relevant saccharification traits, we generated at least 15 models of each type, relying on different fluorescence lifetime parameters, in order to select the best predictive model of each type, as assessed by leave-one-our cross-validation (Dataset S4).

While good correlations could be found between fluorescence lifetime parameters and most cell wall traits (Dataset S4), leave-one-out cross-validation revealed that only two models could yield accurate predictions (Q2 > 0.5): the generalized additive models for G-type and S-type lignin content (Fig. 3C,D; Dataset S4), consistent with the fact that the recorded lignin mostly consists of G and S units in the wood of *Populus* trees. Better predictions for lignin composition were also likely facilitated by the fact that, in addition to using data from our collection of 90 trees, we also included data from lignin samples that either consisted of pure G-lignin or from a synthesized mix of S-type and G-type lignin (Dataset S3).

We leveraged these two models to predict the average S/G-type lignin ratio for the fibres and for the vessels (Fig. 3E,F) of the developing wood samples of a wild-type tree (Fig. 3A) and a tree with reduced expression of the lignin biosynthetic enzyme C4H (*c4h*-RNAi; Fig. 3B). The average S/G-lignin ratio of the fibres was higher than that of vessels in both trees (Fig. 3E,F). The *c4h*-RNAi tree (Fig. 3F), showed a much higher S/G-lignin ratio for both vessels and fibres than in the wild-type tree (Fig. 3E). The overall increase in S/G-type lignin ratio upon downregulation of C4H is also consistent with previous observations^18^ and predictions^51,52^.

We sought to investigate whether FLIM parameters could also help predict the frequency of different covalent bonds between lignin units. To this end, we analysed a subset of ten wood samples by 2D-NMR, which can reveal the frequency of the three most common bonds^53^ between lignin units^54^. These 2D-NMR measurements (Dataset S5) were compared to the average FLIM parameters measured on the corresponding couples of trees (Dataset S5).

Statistical modelling did not reveal any good predictive model of covalent bond types based on fluorescence lifetime parameters (Dataset S4). However, it cannot be excluded that such predictions could be obtained in the future, as the statistical modelling in this case was strongly constrained by the fact that the limited number of 2D-NMR analysis only allowed to generate 10 observations, instead of the at least 90 observations for the aforementioned chemical composition models. Hence, while future correlative analyses between 2D-NMR and FLIM observations of wood cell walls might yield a FLIM readout for covalent bond types in the lignin polymer, at this point we can only confidently predict S-type and G-type lignin content from fluorescence lifetime parameters.

### FLIM on live xylem cells *in vivo* reveals the dynamics of lignin deposition in *Arabidopsis* roots

FLIM should be applicable in live cell imaging as the FLIM instrumentation consists of a laser confocal microscope base. Hence, we set up to image the first root xylem vessel cells that start depositing lignin in the protoxylem cell files of living *Arabidopsis thaliana* seedlings. It had been previously proposed that lignification dynamics would result in G-type lignin being accumulated comparatively earlier than S-type lignin^55-57^. For the first time, this hypothesis can be verified by live cell imaging thanks to our finding that FLIM parameters can be used to estimate S-type and G-type lignin.

In wild-type seedlings, the ratio of S-type to G-type lignin increased within 6h from the first observations (Fig. 4A,B), consistent with previous hypotheses about lignification dynamics (Terashima, 1990; Gerber et al., 2016). In contrast, the *c4h-3* mutants^34^, which display reduced expression of one of the first enzymes in the biosynthesis of lignin monomers (all types), showed a high S/G lignin ratio early, with a decrease after 6h (Fig. 4A,B).

**Figure 4.**
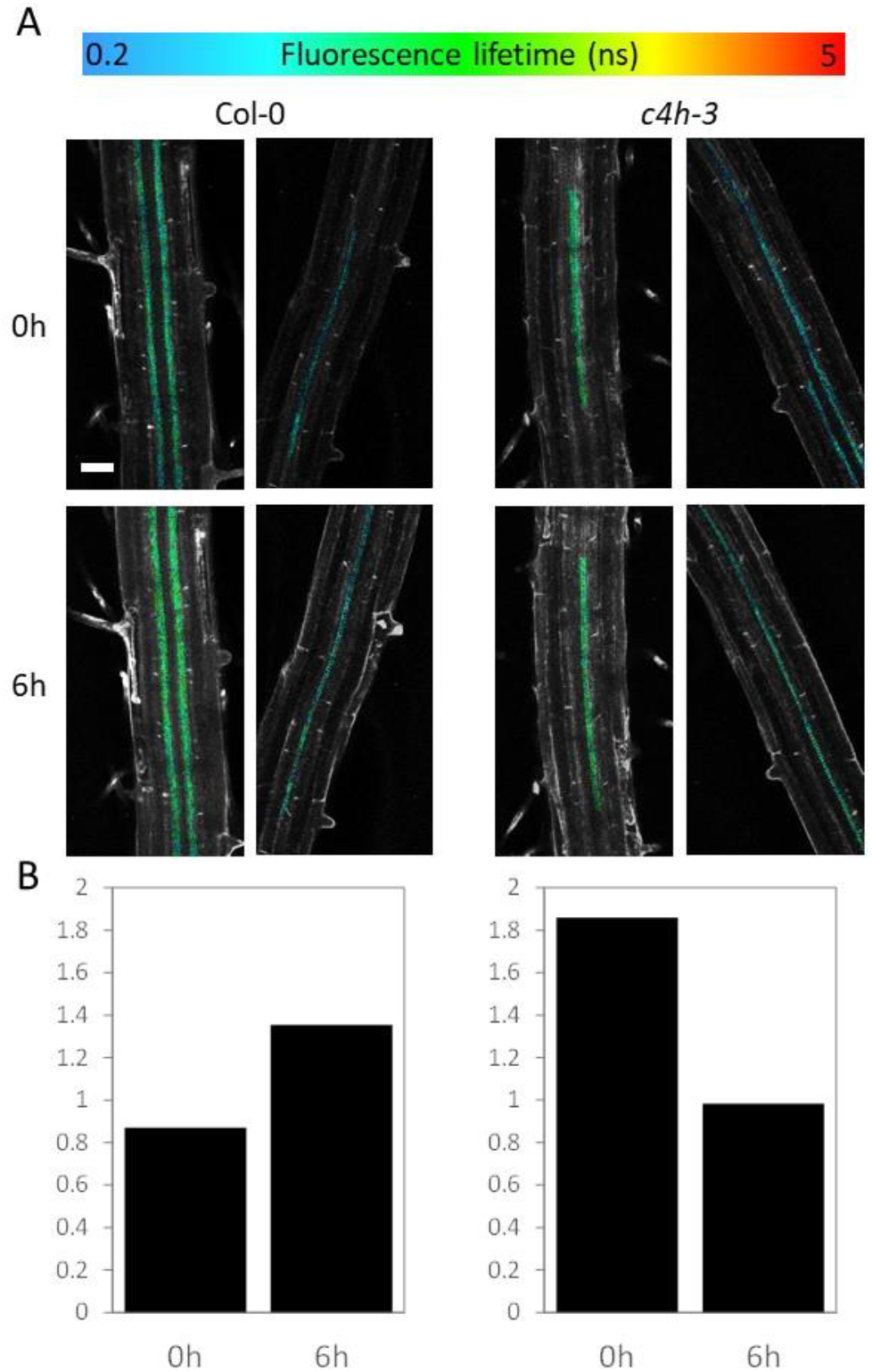
Live imaging of differentiating xylem cells reveals dynamic changes in lignin composition over the course of xylem cell wall biosynthesis. (A) Fluorescence lifetime micrographs of the first lignifying proto-xylem tracheary element cells. The color indicates the average lifetime of the cell wall autofluorescence. Scale bar = 40 µm. (B) Average value of S/G-lignin ratio for each genotype (as in A), as predicted from FLIM parameters using the best statistical models for G-type and S-type lignin (Dataset S4). These average values are from the two different seedlings (in (A)) observed over time in independent experiments.

### FLIM reveals differences in developing cell wall lignin composition in response to tree nutrition

Our use of two fertilization treatments during tree growth allows us to interrogate whether these different fertilizations could lead to differences in wood chemical composition. Based on our mass-spectrometry analyses of homogenized wood, which reflect the average chemical composition across the wood (i.e. not capturing potential local, *in situ* heterogeneity), we could not observe any consistent difference in wood chemical composition matching the different fertilization treatments (Fig. 5A-F). There are, however, notable exceptions to this trend: for two genotypes, the trees having received mineral nitrogen showed higher S-lignin content than the trees having received organic nitrogen (Fig. 5C). For another genotype, higher H-type lignin content was observed under mineral nitrogen nutrition as compared to organic nitrogen nutrition (Fig. 5D). These three genotypes also displayed a different S/G-type lignin ratio under the different types of nutrition (Fig. 5F). For all other genotypes, there was no detectable significant difference in lignin content or composition between mineral and organic nitrogen nutrition (Fig. 5A-F).

**Figure 5.**
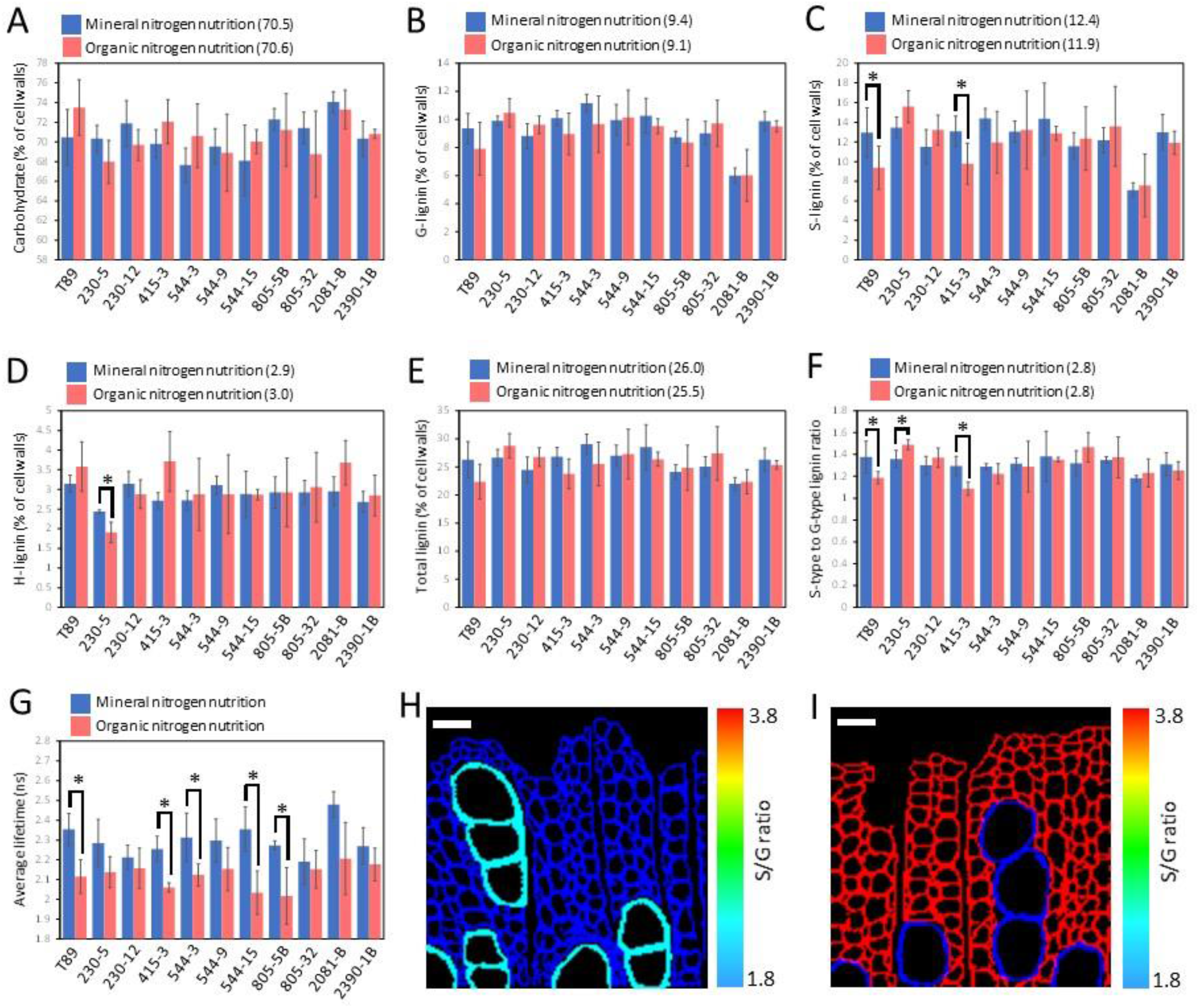
Effect of organic and mineral nitrogen fertilization on lignin chemical composition and lignification dynamics. (A-G) Average value for each genotype under organic (red) versus mineral (blue) nitrogen nutrition, for cell wall contents of carbohydrates (A), G-lignin (B), S-lignin (C), H-lignin (D), total lignin (E), as well as S-type to G-type lignin ratio (F) and cell wall fluorescence lifetime (G). Error bars indicate standard deviation. Asterisks indicate significant differences (p < 0.05) according to a post-ANOVA Fisher LSD test. (H-I) Micrographs of wild-type developing wood under organic (H) or mineral (I) nitrogen nutrition, where the vessels on the one hand, and the fibres on the other hands, were pseudo-colored based on the average S/G-lignin ratio for this cell-type, as predicted in each sample by the statistical models for S-type and G-type lignin content (C, D, and Dataset S4). Average values for S/G-lignin ratio were calculated (for each cell type) from three biological replicates for each nutrition treatment. Scale bar = 30 µm.

In contrast with the average wood chemical composition, the average fluorescence lifetime of the imaged wood section showed a consistent trend across genotypes, with higher fluorescence lifetime values in trees having received mineral nitrogen rather than organic nitrogen (Fig. 5G). This clear trend suggests that while no chemical differences could be detected in homogenized wood samples, there could be greater heterogeneity in wood chemical composition that may be picked up by FLIM.

FLIM observations of specific areas of developing wood, where lignification is ongoing and therefore more dynamic, did reveal differences in lignin composition between wild-type trees treated with organic nitrogen (Fig. 5H) and mineral nitrogen (Fig. 5I). Based on predictions of S-type and G-type lignin content from FLIM images, the cell walls of fibres and vessels had on average a similar S/G-type lignin ratio under organic nitrogen fertilization (Fig. 5H). Under mineral nitrogen fertilization, vessels had on average a much lower S/G-type lignin ratio than fibres (Fig. 5I). The vessels of trees fertilized with mineral nitrogen (Fig. 5I) and both fibres and vessels of trees fertilized with organic nitrogen (Fig. 5H) showed comparable S/G-lignin ratios. Hence, the main difference between the lignin deposited in these fertilization conditions consists in a relative enrichment in S-type lignin (roughly 100 % more) in the fibre cell type upon mineral nitrogen nutrition.

## Discussion

Previous studies had reported that the natural fluorescence of lignocellulosic cell walls showed shifts in fluorescence lifetime either as a result of genetic modification that target lignin biosynthesis^31^, or as a result of chemical treatments altering the chemical and structural environment of lignin^32,33^. Similarly, we observed shifts in fluorescence lifetime when observing a larger sample set of *Arabidopsis thaliana* mutants affected in different aspects of lignocellulosic cell wall composition (Fig. 1A-C). However, these observations also revealed that the relationship between cell wall chemical composition and the average fluorescence lifetime of a sample is not linear (Fig. 1B). This is consistent with the fact that lignin is a complex fluorophore, and implies that several parameters of fluorescence lifetime should be considered in order to read out lignocellulose cell wall chemical composition from FLIM.

Next, we assembled a set of 90 trees from 11 genotypes, selected for their expected changes in cell wall properties, to serve as standard samples to identify relationships that predict lignocellulosic cell wall chemical composition based on FLIM. We observed multiple cell wall composition and saccharification traits (Fig. 2A,B; Dataset S3), as well as multiple parameters of fluorescence lifetime measurements (Fig. 2C; Dataset S3), leading us to find FLIM parameters that could accurately predict either G-type lignin content (Fig. 3C; Dataset S4), or S-type lignin content (Fig. 3D, Dataset S4). The fact that G-type and S-type lignin can be predicted from FLIM is consistent with the fact that lignin is responsible for the fluorescence of lignocellulosic cell walls^22,24^. It is also consistent with the fact that in *Populus* trees, as in other angiosperm species, most of the lignin in the xylem is composed of S-type and G-type units^2^.

A small percentage of lignin can consist of atypical units such as those derived from monolignol biosynthetic intermediates^20,58,59^. Lignin often also comprises a small proportion of conjugated units, whereby classical units bear additional chemical groups^20,60^. Our calibration dataset consisted of measurements where only S-type, G-type, H-type, and “other phenolics” lignin units (Dataset S3) were recorded by our analytical methods, without the much more technically challenging quantification of specific atypical or conjugated lignin units. While it is possible that atypical lignin units would have been accounted for in our analyses as “other phenolics”, it is also possible that the analytical methods we used would miss-classify some atypical or conjugated lignin units as e.g. S-type, or G-type units. These less abundant atypical units may therefore currently be counted as S-type or G-type units in our FLIM-based models. Such hypothetical miss-classification would only affect the FLIM-based measurements of S-type and G-type units by a few percent, which means that the conclusions we draw in this article would remain valid. Yet, future work could use plant genotypes genetically designed to synthesize lignin highly enriched in atypical or conjugated units as standards to help refining our current models, and to enable the detection and quantification of such atypical lignin units by FLIM. The work we present here only lays the foundation for the development of FLIM as a chemical readout on lignocellulosic cell walls, and the possibilities virtually encompass the detection and quantification of any type of lignin units by FLIM in the future.

Other cell wall composition traits or saccharification traits could not be predicted by FLIM in our dataset. The question remains of whether future work could identify a relationship, or if there is no relationship to be found. For example, complex, composite traits such as release of sugars from saccharification might depend on many other traits^18^, only a fraction of which might be related to lignin fluorescence lifetime. In fact, our failure to predict saccharification traits based on FLIM parameters is consistent with the theoretical physics of fluorescence lifetime. The theory indeed implies that fluorescence lifetime is only tied to the chemical composition of the fluorophore^26,28^. Leaning on this theoretical basis, fluorescence lifetime parameters measured on lignocellulosic cell walls should be expected to only relate to the fluorophore lignin, and to other molecules covalently binding to it (e.g. hemicelluloses, consistent with results shown in Fig. 1A-C), but not to more loosely related traits such as sugar release from saccharification.

Based on the same theoretical physics background, FLIM may also display a relationship to the frequency of different covalent bonds between units in the lignin polymer, but we failed to identify such relationships (Dataset S4). A likely explanation resides in the number of observations, as we could only quantify covalent bonds between lignin units in a set of 10 samples (while all other traits could be measured on at least 90 samples). Future studies with a greater number of samples should reveal relationships between FLIM parameters and frequencies of different covalent bonds within lignin. The current study therefore only lays the foundation for using FLIM as an *in situ* readout for lignocellulosic cell wall chemical composition.

Our discovery of models capable of predicting G-type and S-type lignin from FLIM enabled us to monitor the dynamics of lignin deposition in the walls of developing xylem cells *in vivo*, thanks to the fact that FLIM measurements are compatible with live cell imaging. Looking at the first lignifying xylem tracheary element cells in the root of *Arabidopsis thaliana* seedlings, we could visualise a shift in the ratio of S-type to G-type lignin. Our direct, *in vivo* observations revealed that more G-type lignin is deposited early, followed by an enrichment in S-type lignin over the next six hours in wild-type plants (Fig. 4A,B). This is in line with previous, indirect observations which had raised the question as to whether G-lignin would be mostly deposited earlier than S-lignin^55-57^.

The dynamics of lignification were different between wild-type plants and the *c4h-3* mutants genotypes (Fig. 4A,B), whose mutation is known to affect lignin monomer biosynthesis in the cytoplasm. Observing differences in the dynamics of lignification between these genotypes is therefore in line with the idea that lignin composition in xylem cell walls mostly results from random polymerization, and is therefore largely determined by the dynamics of monomer biosynthesis in the cytoplasm^2^.

The 11 tree genotypes used in this study were grown under two different nutrition conditions: nutrition with a nitrogen source mostly in an organic form, and a more classical nutrition with nitrogen mostly in the mineral form (Dataset S2). These different treatments did not result in consistent changes in wood chemical composition across the entire tree population, although specific genotypes showed significant differences in accumulation of S-type or H-type lignin (Fig. 5C,D). Wild-type trees in particular showed increased S-type lignin content and increased S-type to G-type lignin ratio under mineral nutrition treatment. Using FLIM, we observed the developing wood of wild-type trees and also found an increase in S-type to G-type lignin ratio under mineral nutrition (Fig. 5H,I). These FLIM observations *in situ* also revealed that these differences in lignin between treatment occurred specifically in the xylem fibre cell type, but not in vessels (Fig. 5H,I), providing an interesting anatomical resolution to the analysis of wood chemical composition between treatments.

Interestingly, our observations of developing wild-type wood with FLIM showed even greater differences in S-type to G-type lignin ratio between nutrition treatments (Fig. 5H,I) than those observed on homogenized wood powder (Fig. 5F). Such quantitative differences may arise from lignification dynamics whereby early lignification of wood under mineral nutrition would consist in an overwhelming majority of S-type lignin in fibres, followed by a later relative enrichment in G-type units. Under organic nutrition, the dynamics would be opposite, with an early accumulation of G-type lignin, followed by enrichment in S-type lignin in fibre cell walls over the course of wood formation.

FLIM has previously been applied to detecting chemical differences in biological samples^26^, for purposes of classification such as distinguishing pathological versus healthy samples in diagnosis^61^, or such as distinguishing between metabolic states or between differentiation states in living cells^62^. These strategies, however, do not quantify aspects of the chemical composition of the fluorophores based on FLIM. Our approach using FLIM for chemical characterization of naturally fluorescent lignocellulosic cell walls shows that it is possible to take a significant step further: we provide an example that FLIM can be used for *in situ* quantitative measurements of the chemical composition of naturally fluorescent molecules, even *in vivo*. Similar to lignin, other important biological molecules are known for their fluorescent properties. Such molecules include collagen^63^, keratin and elastin^61^, NADH^62^, or resillin in the shell of insects^64^. These molecules could in theory also be chemically characterized *in situ* by FLIM, if similar efforts were undertaken for the calibration of statistical models as those we present here for plant lignocellulosic cell walls.

## Materials and Methods

### Plant material and growth conditions

The *Arabidopsis thaliana* genotypes consisted of Col-0 wild-type; the *c4h-3/ref3-3* which contains a point mutation in the cinnamate-4-hydroxylase enzyme-encoding gene (*C4H*; At2g30490) that results in a still partially functional protein^34^; the *gux1 gux2* double mutant previously generated by crossing of the T-DNA lines SALK_046841 (AT3G18660) and GABI_722F09 (AT4G33330) that lack expression of functional glucuronic acid substitution of xylan (GUX) proteins^36,37^; and the irx7/fra8 mutant which harbours a point mutation in a gene (At2g28110) encoding a glycosyltransferase family 47 protein involved in xylan biosynthesis^39^. For imaging of stem cross-sections, these plants were grown for eight weeks in growth chambers with 16h simulated day-light and 8h night, with semi-controlled temperatures oscillating between 18°C and 23°C. After these eight weeks, a 1 cm long piece of stem was sampled 2 cm over the soil for further sectioning and FLIM observations. For live imaging by FLIM of the first lignifying protoxylem cells, 4.5 days old (after germination) seedlings of Col-0 and *c4h-3* were used.

The collection of hybrid aspen (*Populus tremula* x *tremuloides* Michx.) trees used as “standards” to relate wood chemical composition to FLIM parameters consisted of 11 genotypes: the wild-type T89 clone, as well as ten transgenic lines, targeting six genes known or thought to be linked to wood cell wall properties (Dataset S1^18,65-68^). To generate replicates, the trees were clonally propagated *in vitro* as previously described^69^. After three weeks of *in vitro* growth, eight (all transgenics) or ten (wild-type T89) trees from each genotype were transferred to 1 L soil pots and grown for nine weeks under 16h/8h day-night cycles. Following an initial period of three weeks of tree establishment in the soil conditions, the trees started receiving fertilization with increasing doses of either one of two nutrient solutions (Dataset S2): the arGrow solution (Arevo AB, Sweden) with nitrogen mostly in the organic form arginine, and SW Bouyant Rika-S (SW Horto AB, Sweden) with nitrogen mostly in the mineral form of nitrate (Dataset S2).

### Specific lignin samples

The three samples of extracted pine lignin were prepared by dioxane/HCl acidolysis following the “reflux method”^70^, but with some modifications: pine wood samples were milled to a particle size of 200 µm (Retsch ZM1000 centrifugal mill, Retsch GmbH, Germany). 20 g of the milled samples were placed in 250 mL dioxane/HCl 2 M (9:1 v/v) under a flow of nitrogen gas for 3 h. Another 250 mL of dioxane/HCl 2M were added and the solution was heated to a reflux for 30 min. This solution was filtered and poured onto 8.4 g of NaHCO3 for acid neutralisation. Following the near complete evaporation of the dioxane, the samples were thoroughly washed with large volumes of ultrapure water, centrifuged and finally air-dried.

The synthesized lignin samples, so called dehydrogenation polymers (DHP), were generated *in vitro* as previously described^71,72^, but with an equal mix of coniferyl alcohol and sinapyl alcohol to generate a polymer of S-type and G-type lignin units with a 1:1 ratio.

### Sampling and preparation for different analyses

After nine weeks of growth, the 90 trees were harvested (Fig. S2). Stem pieces were collected 15 cm above-ground to generate the samples for all the cell wall chemistry and imaging analyses. From these stem pieces, several sections were made: The 9 cm towards the base of the stem were freeze-dried (CoolSafe Pro 110-4, LaboGene A/S, Denmark), milled (Retsch ZM200 centrifugal mill), and the resulting wood powder with particle size of less than 0.5 mm was collected for subsequent 2-dimensional nuclear magnetic resonance (2D-NMR) analyses. The next 1 cm was kept apart, and the following 2 cm of stem were collected and 100 µm sections were generated using a vibratome (Leica VT1000S, Leica Biosystems GmbH, Germany). These sections were mounted in 0.1 M phosphate buffer at pH 5.8 (close to the physiological pH of plant cell walls) for direct observation by FLIM. The next 4 cm of stem were freeze-dried, milled, and sieved to separate particles of different sizes. Particles with a diameter between 0.1 mm and 0.5 mm were collected for enzymatic saccharification assay. Particles with diameter lower than 0.1 mm were collected separately for analysis of cell wall lignin content and composition by pyrolysis coupled with gas chromatography and mass spectrometry.

*Arabidopsis thaliana* stem samples, collected 2 cm above-ground after eight weeks of plant growth, were cut with a vibratome as described above to generate 100 µm sections. These sections were also mounted in 0.1 M phosphate buffer at pH 5.8 and observed by FLIM soon after.

### Analyses of wood cell wall chemical composition and saccharification

The cell wall content in carbohydrates (indiscriminately cellulose, hemicelluloses, and other possible saccharides), as well as the lignin content and composition, were all measured by pyrolysis gas chromatography coupled with mass spectrometry (pyrolysis-GC/MS), as previously described^56^.

Enzymatic saccharification assays, measuring the release of five monosaccharides (glucose xylose, galactose, mannose and arabinose) either without pretreatment or after an acidic pretreatment, were performed on the wood powder of particle size 0.1 mm to 0.5 mm, as previously described^73^ using a liquid enzyme preparation (5 mg Cellic CTec2, obtained from Sigma-Aldrich, St. Louis, MO, USA). The concentrations of the five monosaccharides were determined using HPAEC-PAD (ICS-5000, Dionex, Sunnyvale, CA, USA) following a previously described procedure^74^. The glucose production rate (GPR), which is based on the initial two hours of saccharification, was determined as previously described^75^.

Analysis of the relative abundance of covalent bonds within the lignin polymers relied 2D-NMR, as previously described^54^. These analyses required pooling of samples to gather a sufficient amount of wood powder with particle size between 0.1 mm and 0.5 mm, and could only be performed on a subset of trees (Dataset S5).

### Imaging and image analysis

Fluorescence Lifetime Imaging Microscopy images were acquired directly on the sections of tree stems or *Arabidopsis thaliana* stems, solely based on lignin natural fluorescence. All fluorescence lifetime imaging microscopy observations relied on the time-correlated single photon counting methodology (TCSPC; Becker, 2005).

The *Arabidopsis thaliana* stem sections were imaged with a confocal microscope LSM710 (Carl Zeiss Microscopy GmbH, Germany) using a Chameleon TiSa accordable 80 Mhz pulsed laser (Coherent Inc., USA) for excitation, and a MW-FLIM detector system with a SPC 150 photocounting card (Becker & Hickl GmbH, Germany) for detection, as previously described^32^. Images were processed using SPCImage 6.2 (Becker & Hickl GmbH, Germany) to extract fluorescence lifetime data specifically from xylem cell walls (Fig. 1), selected as regions of interest (ROIs).

The synthesized and extracted lignin samples, as well as the tree stem sections, were imaged with a confocal microscope LSM880 (Carl Zeiss Microscopy GmbH, Germany) coupled with a FLIM Upgrade Kit for LSMs (PicoQuant GmbH, Germany), including PicoQuant pulsed diode lasers (tuneable laser repetition rate between 2,5 MHz and 40 MHz) and HyD single molecule detectors (SMD) for photon detection and counting. The aim being to correlate FLIM parameters to measurements from other analytical methods that are conducted on homogenized stem material, FLIM images needed to be acquired for the entire wood surface considered. Due to the large areas to be imaged, tile imaging had to be performed (4 × 4 tiles with 10% overlap) with a x4 lens when looking at entire sections. In addition, images of developing wood areas were also acquired with a x20 lens for higher resolution.

Images were processed, and the corresponding FLIM parameters were extracted using the SymPhoTime64 software (SPT64, PicoQuant GmbH, Germany). FLIM fitting on the lifetime decay curves revealed that the best fits could be obtained using three exponentials, meaning that the average lifetime could be best described as a composite resulting from three different lifetimes. FLIM parameters (Dataset S3) were extracted on whole sections, allowing comparison with the analytical methods that record cell wall chemical composition of homogenized ground samples (Dataset S3). FLIM parameters could also be extracted from specific wood areas, or specific cell types, by using the region of interest (ROI) function of the SPT software, allowing for manual selection of any pixel or group of pixels on the images.

### Statistical modelling

Statistical modelling followed a previously established methodology^18,43^ that consists in using three different types of models to select the most predictive model based on leave-one-out cross validation. Briefly, statistical relationships between FLIM parameters (independent variables) and each wood cell wall or saccharification trait (dependent variable) was explored by multiple linear regression, as well as by the more complex machine learning algorithms known as generalized additive models (GAMs)^49^ and random forests^50^.

All types of models were implemented in R, using the package “mgcv” for GAMs^76^ and the package “ranger” for random forests^77^. For each dependent variable, at least 15 models from each type (i.e. at least 45 models) were generated. In addition to the variation explained by each model (R2), we assessed whether a model was suitable for prediction by leave-one-out cross validation. This validation consists in leaving out one observation (one tree) out of the dataset, re-generating the model based on the remaining observations (the 89 other trees), and predicting the value of the one tree that had been left out. This procedure is repeated until each observation had been left out, and the difference between the predictions and the actual values for each tree allow calculating the Q2 (with Q2 between 1 and minus infinity, 1 indicating a perfect prediction).

## Author contributions

SE and GP designed the study, with conceptual contributions and feedback from CT, GS, TNä, OS, LJJ, JLF, HT, and TNi. SE, TNä, JLF, and HT contributed with the design and production of genetically engineered trees used in this study. SE, CT, MLG and ZY performed the experiments: SE grew the plants, harvested the samples, and prepared the stem sections. CT and SE performed FLIM imaging of *Arabidopsis thaliana* stems. SE performed FLIM imaging of wood stems, synthesized and extracted lignins, as well as live imaging of lignin deposition dynamics in *Arabidopsis thaliana* seedlings. SE, MLG and ZY prepared the samples for all other analytical methods. MLG performed the saccharification assays (including pretreatments when relevant). SE extracted data (lifetime parameters) from FLIM analyses, and performed statistical analyses and modelling. SE and GP wrote the manuscript, with help from all co-authors.

## Acknowledgements

The authors wish to acknowledge the valuable technical help from Junko Takahashi-Schmidt (pyrolysis-GC/MS, UPSC biopolymer analytical platform), Anouck Habrant (synthesis of DHP, FARE lab), and David Crônier (2D NMR analysis, FARE lab). This work was funded by the Swedish Strategic Research Environment Bio4Energy (https://www.bio4energy.se) through two grants: the Bio4Energy Special Call Grant 2018, and the Bio4Energy Strategic Funds Grant 2019. Bio4Energy is one of three strategic research environments funded by the Swedish Energy Agency.

